# Nitrogen starvation induces persister cell formation in *Escherichia coli*

**DOI:** 10.1101/445957

**Authors:** Daniel R. Brown

## Abstract

To cope with fluctuations in their environment bacteria have evolved multiple adaptive stress responses. One such response is the nitrogen regulation stress response, which allows bacteria such as *Escherichia coli* to cope with and overcome conditions of nitrogen limitation. This response is directed by the two-component system NtrBC, where NtrC acts as the major transcriptional regulator to activate the expression of genes to mount the response. Recently we showed that NtrC directly regulates the expression of the *relA* gene, the major (p)ppGpp synthetase in *E. coli*, coupling the nitrogen regulation stress and stringent responses. As elevated levels of (p)ppGpp have been implicated in the formation persister cells, here we investigated whether nitrogen starvation promotes their formation and whether the NtrC-RelA regulatory cascade plays a role in this. The results reveal that both nitrogen starved *E. coli* form a higher percentage of persister cells than non-starved cells, and that both NtrC and RelA are important for this process. This provides novel insights into how the formation of persisters can be promoted in response to a nutritional stress.

**Importance:** Bacteria often reside in environments where nutrient availability is scarce and therefore they have evolved adaptive responses to rapidly cope with conditions of feast and famine. Understanding the mechanisms that underpin the regulation of how bacteria cope with this stress is a fundamentally important question in the wider context of understanding the biology of the bacterial cell and bacterial pathogenesis. Two major adaptive mechanisms to cope with starvation are the nitrogen regulation (ntr) stress and stringent responses. Here I describe how these bacterial stress responses are coordinated under conditions of nitrogen starvation to promote the formation of antibiotic tolerant persister cells by elevating levels of the secondary messenger (p)ppGpp.

## Introduction

The survival and proliferation of bacteria requires them to adapt to fluctuating availability of macro- and micronutrients. To overcome these challenges, bacteria have evolved strategies in the form of adaptive stress responses to sustain growth on non-optimal sources of nutrients, outcompete others for available resources or enter a state of metabolic quiescence to survive until conditions improve. One such response is the nitrogen regulation (ntr) stress response, which allows *E. coli* and related bacteria to cope with conditions of nitrogen starvation (1, 2). Nitrogen (N) is an essential element for life, as numerous cellular macromolecules contain nitrogen, including proteins, nucleic acids and cell wall components. This adaptive response is coordinated by the two-component system NtrBC. Where NtrB is the sensor histidine kinase that senses and then transduces the nitrogen starvation signal upon which it phosphorylates and thereby activates the DNA binding transcription factor NtrC (3). NtrC is a member of a specialised family of transcriptional activators named the bacterial enhancer binding proteins (bEBPs), which activate the alternative transcription machinery directed by the alternative sigma factor σ^54^ in response to diverse environmental stimuli (4, 5). The ntr response allows cells to rapidly sense and adapt to nitrogen limitation. It does this by actively scavenging for alternative nitrogen sources through the transcriptional activation of genes encoding nitrogen source transporters, catabolic enzymes and amino acid biosynthetic operons (1). The response also induces further expression of genes to survive N starvation through the action of the dual regulator Nac, the nitrogen accessory control protein (6). Recently we showed that the least understood NtrC-regulated operon, *yeaGH*, which encodes a putative serine/threonine kinase and a gene of unknown function respectively, promotes survival of *E. coli* faced with sustained N starvation, appearing to act as a molecular brake, dampening metabolic activity of N starved *E. coli* (7, 8).

In a previous study we sought to determine the NtrC regulon in N-starved *E. coli.* This identified all of the above genes as being under the direct transcriptional control of NtrC (9). Interestingly, this genome-wide analysis also revealed a novel member of the NtrC regulon and uncovered a new branch to the ntr response. We discovered that NtrC binds upstream of, and activates, the σ^54^-dependent transcription of the *relA* gene, which encodes for the major (p)ppGpp synthetase in *E. coli* (9). Increased intracellular levels of the secondary messengers (p)ppGpp have pleiotropic effects upon major cellular processes such as transcription, translation and DNA replication to reprogram the cell to cope with the change in the environment, in particular with regard to nutrient starvation (10). This action of (p)ppGpp is collectively known as the stringent response. Our finding illustrated an elegant mechanism in which nitrogen starved *E. coli* use the same regulatory mechanism to concurrently activate the transcription of genes to actively scavenge for alternative sources of nitrogen to survive whilst switching off genes required for exponential growth. The induction of (p)ppGpp has been previously implicated with the increase in levels of so-called persister cells (11-14), which are a small sub-population of bacteria which can transiently tolerate high levels of antibiotics without being genetically resistant (11, 14). The formation of persisters is induced under a variety of conditions, including during infection (12). They have also been proposed as a reservoir for recurrent infection and are potentially a basis for development of antibiotic resistance (15). Therefore, understanding the regulation that underpins their formation, maintenance and resurgence are key questions which have yet to be fully answered. Previous work has shed light upon a multitude of conditions to understand how persisters form, such as different growth phases in rich medium, growth on defined carbon sources, and even the host environment (macrophages) (12, 16-19). In our case, we wanted to investigate whether nitrogen starvation induced the formation of persister cells, and whether NtrC-dependent gene expression, in particular that of *relA*, plays a role in this.

## Results and Discussion

### Nitrogen starvation induces an increase in ciprofloxacin-tolerant persisters

The initial aim of this study was to understand whether *E. coli* subjected to N starvation form elevated levels of antibiotic tolerant persisters in comparison to nonstressed bacteria. To accomplish this, we used the well-defined nitrogen starvation conditions we previously established to study the nitrogen stress response (9, 20). To ensure that cells in the non-stressed condition (N+) were not sensing limitation to nitrogen, and therefore to ensure they were in a “non-stressed state” they were grown in Gutnick minimal medium containing 10mM NH4Cl prior to exposure to antibiotics at early log phase. Nitrogen-starved cells (N-) were exposed to antibiotics 20 minutes post runout of nitrogen following growth in Gutnick minimal medium containing 3mM NH_4_Cl. Growth rates of the cells are comparable when grown in either 10mM or 3mM NH_4_Cl, as show in Fig. 1a. As can be clearly seen, addition of antibiotics (>10x MIC, ciprofloxacin MIC’s of all strains used in this study are the same and can be seen in Table 1.) at either of these time-points (N+ or N-) results in characteristic biphasic kill curves, with an initial fast killing of the susceptible population which then slows over time (Fig. 1b). When comparing the two conditions, there is a ~20-fold increase in survival of the N starved (N-) wild-type cells compared to non-starved (N+) cells following 24 hours of exposure to ciprofloxacin (Fig. 1c). Thus, indicating that there is a higher proportion of persisters in the bacterial population subjected to N starved conditions compared with the non-stressed bacterial population.

**Figure 1.**
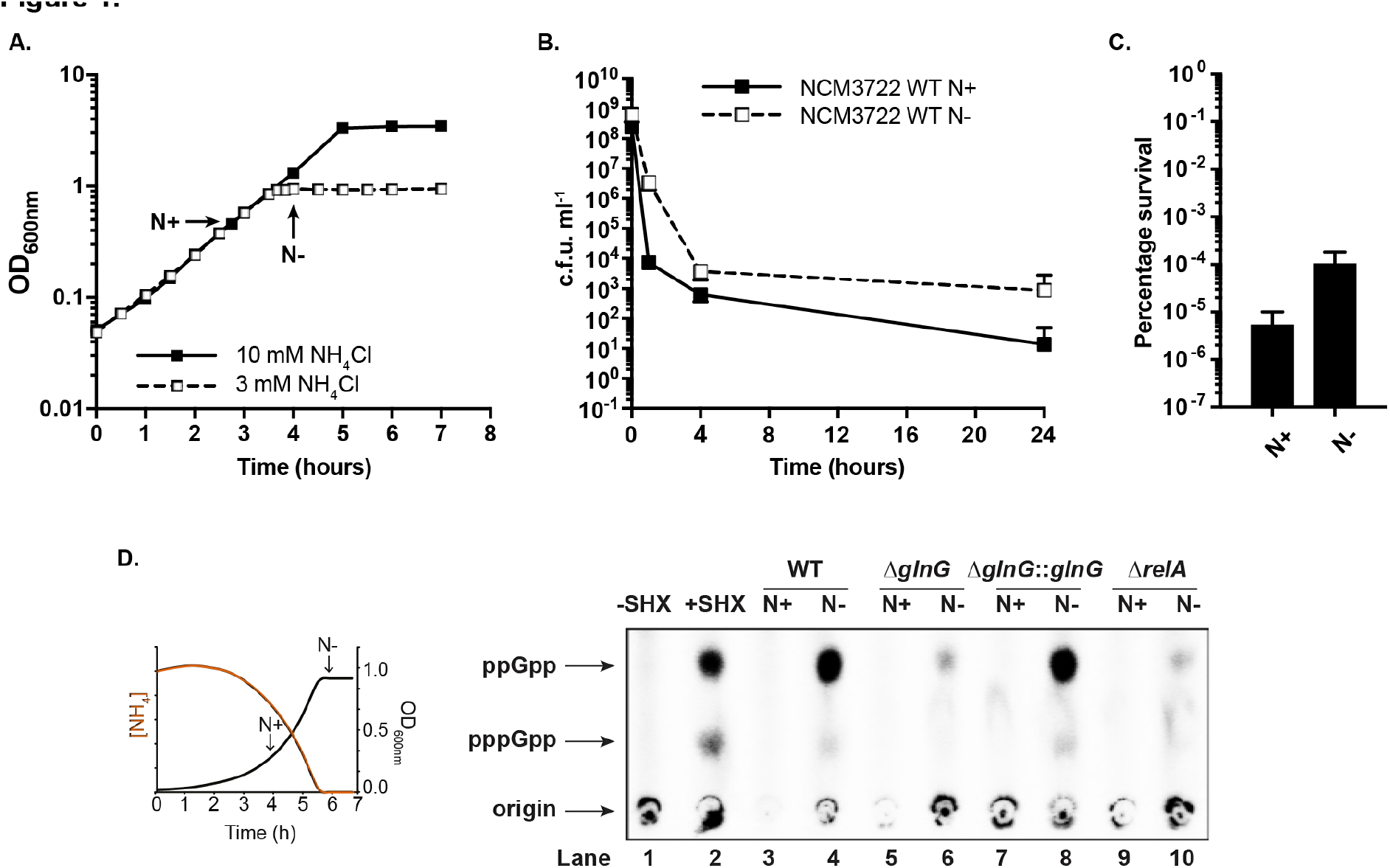
Nitrogen starvation promotes ciprofloxacin-tolerant persister formation. **(A.)** Representative growth curve of NCM3722 wild-type *E. coli* in Gutnick minimal medium containing 3mM (dashed line) or 10mM (solid line) NH_4_Cl as the sole N source. Time-points used for persister assays are highlighted (N+ = nitrogen replete & N- = nitrogen starved). **(B.)** Persister assay results displayed as c.f.u. ml^-1^ to show bacterial survival following N+ (solid line) or N- (dashed line) challenged with 1μg/ml ciprofloxacin for 24 hours. **(C.)** Persister assay results displayed as percentage survival of bacteria at 24 hours with respect to pre-antibiotic treatment. Error bars indicate standard deviation from the mean (n=6). **(D.)** Autoradiogram to show levels of (p)ppGpp from ^32^P labelled nucleotide extracts at N+ and N- in NCM3722 strains separated by PEI-cellulose thin layer chromotograpy. Control lanes 1 and 2 show levels of (p)ppGpp in (p)ppGpp non-inducing (-SHX) and inducing (+SHX) conditions. (inset) schematic to indicate growth conditions and time-points assayed for (p)ppGpp measurements.

**Table 1.** Minimum inhibitory concentration of ciprofloxacin measured using MIC Evaluator test

| Strain | Grown in LB | Grown in Gutnick |
| --- | --- | --- |
| NCM3722 wild type | 0.015µg/ml | 0.015µg/ml |
| NCM3722 $\Delta glnG$ | 0.015µg/ml | 0.015µg/ml |
| NCM3722 $\Delta glnG::glnG_{ind}$ | 0.015µg/ml | 0.015µg/ml |
| NCM3722 $\Delta nac$ | 0.015µg/ml | 0.015µg/ml |
| NCM3722 $\Delta rpoS$ | 0.015µg/ml | 0.015µg/ml |
| NCM3722 $\Delta relA$ | 0.015µg/ml | 0.015µg/ml |

In a previous study we found that NtrC, the master regulator of the ntr response regulated the transcription of *relA* in N-starved *E. coli* (9). To show that N starvation leads to the production of RelA synthesized (p)ppGpp here we measured (p)ppGpp and levels at nitrogen replete (N+) and nitrogen starved (N-) conditions. As can clearly be seen in Figure 1.d. (p)ppGpp is produced by the wild type strain following N starvation, but not under N replete conditions (Fig 1.d. lanes 3 and 4). This is also seen in the control lane (Fig 1.d. lanes 1 & 2) when bacteria are subjected to SHX treatment which is known to induce (p)ppGpp production (11). It is also clear that (p)ppGpp synthesis in N starved *E. coli* is dependent upon the action of NtrC (encoded by *glnG*), as in the Δ*glnG* mutant (p)ppGpp is not induced (Fig 1.d. lane 6). Further reinforcing this (p)ppGpp synthesis is restored in the N starved complemented mutant expressing *glnG* from a plasmid (Fig 1.d. lane 8). The (p)ppGpp produced under N starvation conditions appears to be mostly derived from the (p)ppGpp synthetase RelA as in a Δ*relA* mutant very little (p)ppGpp can be detected at either timepoint (Fig 1.d. lanes 9 & 10). This reveals that (p)ppGpp synthesis in N starved *E. coli* is regulated in an NtrC and RelA dependent manner.

### NtrC-dependent gene expression promotes persister formation in N starved *E. coli*

To understand whether persisters formed under N starvation require the action NtrC next I determined levels of persisters formed in its mutant, Δ*glnG*, in the N starvation growth conditions. Strikingly, despite seemingly similar persister levels to the wild-type under non-stressed conditions (N+) a mutant strain lacking NtrC (Δ*glnG*) survives much worse in comparison to the wild-type when challenged with ciprofloxacin following N starvation, ~260 fold fewer persisters are formed (Fig. 2a). Importantly, this phenotype is restored when the *glnG* mutation is complemented via expression of an inducible copy of *glnG* from a plasmid (Fig. 2b). This indicates that under N starvation NtrC-dependent gene expression is important for the formation of ciprofloxacin-tolerant persister cells.

**Figure 2.**
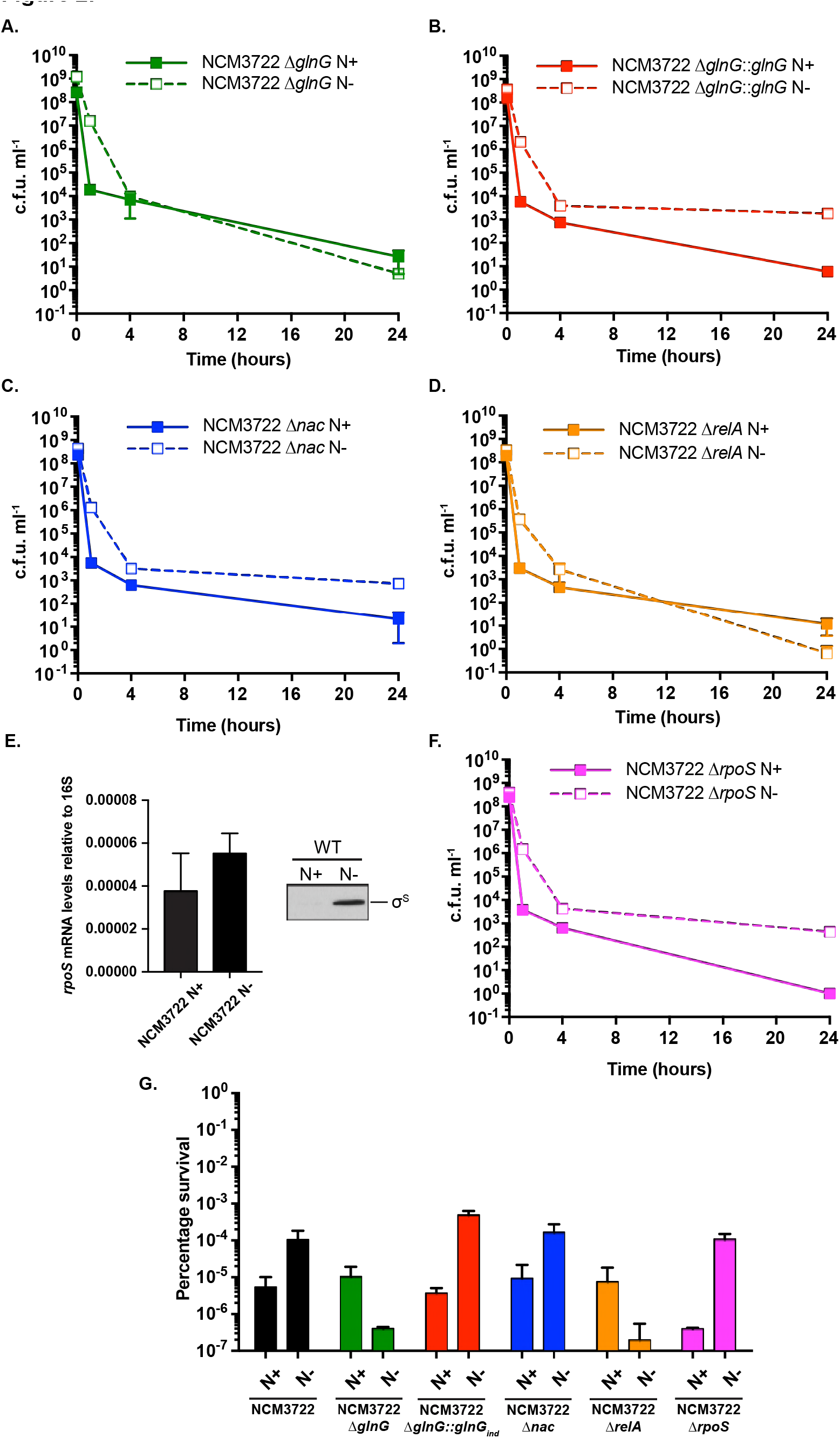
NtrC and RelA are required for the formation of ciprofloxacin-tolerant persisters formed in response to nitrogen starvation. **(A.-D. and F.)** Persister assay results displayed as c.f.u. ml^-1^ to show bacterial survival following N+ (solid line) or N- (dashed line) challenged with 1μg/ml ciprofloxacin for 24 hours. Strain names are inset on each graph **(E.)** qRT-PCR and western blotting analysis to show *rpoS* mRNA and σ^S^ protein levels at N+ and N- **(G.)** Persister assay results displayed as percentage survival of bacteria at 24 hours with respect to pre-antibiotic treatment. Error bars indicate standard deviation from the mean (n=3 for mutants 6 for wild-type).

To investigate whether the regulatory pathway that promoted persister formation under N starvation was via genes whose transcription is activated directly by NtrC (σ^54^-dependent) or by genes regulated by the NtrC-adapter protein Nac (σ^54^-independent), we next determined the ability of a Δ*nac* mutant to form persisters under both N+ and N- conditions. Figs. 2c. & 2f. show that following 24 hours of exposure to ciprofloxacin the N starved Δ*nac* mutant forms similar levels of ciprofloxacin tolerant persisters to the wild-type strain. This indicates that Nac-regulated genes do not impact persister formation, and therefore genes whose transcription is directly activated by NtrC and σ^54^ are the cause for this phenotype. The σ^54^ mutant does not grow in this minimal medium (please see supplementary Fig. 1. which shows the growth of each of the strains used in this study under the experimental conditions), therefore the strict reliance on σ^54^ could not be tested, however, as NtrC can only activate transcription utilizing the σ^54^-RNAP (4, 5) we are confident that σ^54^-dependent transcription is involved in this regulatory process.

### NtrC activated *relA* is required under N starvation to promote persister formation

As in our previous study we found that NtrC activates the expression of the *relA* gene in N starved *E. coli*, recall that RelA is the major (p)ppGpp synthase under these conditions (Fig. 1.d. and Brown et al. 2014), and that (p)ppGpp has been implicated in formation of persister cells, it was logical to determine whether the *relA* mutant strain could form persisters following N starvation. Results in Fig. 2d. clearly show that the Δ*relA* strain is impaired in persister formation in N starved *E. coli*. This implicates the NtrC-relA regulatory cascade in persister formation.

Previous studies have indicated that (p)pGpp plays a role in persister formation (11-14). It is also known that an increase in (p)ppGpp levels correlate with increased stability of the general stress σ factor, σ^38^ or RpoS, and therefore increased levels of RpoS and the gene expression of its regulon (21). RpoS has also been implicated in persister formation, with either mutants lacking *rpoS* or affecting RpoS stability producing both fewer or greater levels of persisters depending on the experimental setup (22-26). Although the gene expression programme most associated with nitrogen stress is directed by the Ntr response and therefore NtrC, σ^54^ and Nac, RpoS-dependent gene expression may also play a role under these conditions. As can be seen in Fig. 3.e. RpoS is clearly upregulated, and RpoS-dependent gene expression has been demonstrated to be active under our N starvation conditions (8). Therefore, we decided to determine whether RpoS plays a role in the formation of N starvation induced persisters. However, as can be seen in Fig. 3.f. the N starved Δ*rpoS* mutant forms similar levels of persisters to the wild-type strain. Therefore, ruling out its role in promoting the formation of persisters under these conditions.

## Conclusions

Determining the mechanisms by which bacteria tolerate stress are fundamental in understanding how they live such adaptable lifestyles and manage to occupy such diverse niches. Despite extensive studies to understand what conditions induce the formation, maintenance and reawakening of antibiotic tolerant persister cells the regulatory factors that underpin their formation in response to stressful environments are not fully understood. Previous studies have however have shown that an increase in (p)ppGpp levels lead to the formation of higher levels of persister cells (11, 13). Results presented here show that N starved *E. coli* synthesise (p)ppGpp and form elevated levels of ciprofloxacin tolerant persisters. The action of NtrC and RelA are key for both of these processes, as neither the Δ*glnG* nor the Δ*relA* mutants synthesize significant (p)ppGpp or generate persisters in response to N starvation. It is interesting to note however, that RpoS which has been implicated with the formation of persisters, is stabilised through the action of (p)ppGpp and is present under N starvation is not required for persisters to be formed under these conditions. Overall these results display an apparent cross-talk between regulatory mechanisms to survive a specific stress, N starvation, and those which allow bacteria to survive high-doses of antibiotics. This further underlines the important role that the alarmone (p)ppGpp plays in stress tolerance of bacteria. This study supports the hypothesis that (p)ppGpp is a key player in persister formation, however the precise mechanism through which it does so remains elusive.

## Methods and materials

### Bacterial strains, plasmids and growth conditions

All strains used in this study were derived from *E. coli* NCM3722, a derivative of *E. coli* K-12 strain MG1655. The NCM3722:Δ*glnG* and Δ*relA* strains were constructed as described previously (9, 20); briefly, the knockout *glnG* allele was transduced using the P1vir bacteriophage with JW3839 for *glnG* and JW2755 for *relA* from the Keio collection (27) serving as the donor strain and NCM3722 as the recipient strain. The kanamycin cassette was then cured from the strain using pCP20 leaving an unmarked knockout mutation. The NCM3722 Δ*rpoS* and Δ*nac* strains were constructed in the same way with E. coli JW5437 and JW1967 (Keio collection) respectively, serving as the donor strain (27). A glnG-inducible complementing plasmid (pAPT-glnG-rbs31) was kindly provided by Wang et al (28). Bacteria were grown in Gutnick media (33.8 mM KH_2_PO_4_, 77.5 mM K_2_HPO_4_, 5.74 mM K_2_SO_4_, 0.41 mM MgSO_4_), supplemented with Ho-LE trace elements (29) and 0.4% glucose, and containing either 10 mM NH_4_Cl (for precultures and N+ persister assays) or 3 mM NH_4_Cl (runout experiments and N- persister assays) as the sole nitrogen source at 37°C, 180 r.p.m. Ammonium concentrations in the media were determined using the Aquaquant ammonium quantification kit (Merck Millipore, UK), according to the manufacturer’s instructions.

### Antibodies

For immunoblotting, the following commercial mouse monoclonal antibodies were used as a primary antibody, anti-σ^S^ (at a dilution of 1/500) (Neoclone, USA). The primary antibody was used in conjunction with the anti-mouse ECL horseradish peroxidase (HRP)-linked secondary antibody (at a dilution of 1/10,000) and ECL prime (GE Healthcare, UK).

### (p)ppGpp measurements

(p)ppGpp levels were measured as described by Schneider et al (30) with the following modifications. Overnight cultures were prepared in 5 ml potassium morpholinopropane sulfonate (MOPS) minimal media containing 10mM NH_4_Cl as the sole nitrogen source. Following growth at 37°C shaking at 180 rpm for 18hours these pre-cultures were used to inoculate 5 ml MOPS minimal media containing 3mM NH_4_Cl as the sole nitrogen source at an OD_600nm_ = 0.05 in 50ml conical tubes. Bacteria were grown at 37°C shaking at 180rpm, bacterial growth was monitored by OD600nm readings. At OD_600nm_ = 0.1, 20 μCi/ml of [^32^P] H_3_PO_4_ was added as a phosphate source. At specified timepoints (N replete when OD_600nm_=0.3 (N+), and 20 minutes post N run out (N-)) 500μl samples were removed and added directly to 100 μl ice cold 2 M formic acid, samples were incubated on ice for 30 minutes prior to centrifugation at 17,000 x *g* for 5 minutes. To detect (p)ppGpp production 10 μl of supernatant from each sample was spotted on a polyethylenimine cellulose F (dimensions 20 cm × 20 cm; Merck Millipore) TLC plate. The spots were then allowed to migrate to the top of the sheet (~70 minutes) in a TLC tank using 1.5M KH2PO4 pH 3.6 running buffer. The plate was then air dried before exposure onto a phosphor screen, following which it was viewed on using a Typhoon Phosphorimager (GE Healthcare). As a positive control bacterial cultures were grown in the same way however 100 μg/ml serine hydroxymate was added at OD_600nm_ = 0.45 to induce (p)ppGpp production, incubated for 10 minutes and samples recovered in the same way as described above.

### Persister assays

Persister assays were completed as described in (12) with the following modifications: Bacteria were routinely grown as 50 ml cultures in either N+ (Gutnick minimal media containing 10mM NH_4_Cl) or N- (Gutnick minimal media containing 3mM NH_4_Cl) in 250ml Erlenmeyer flasks, growth being monitored via optical density measurements. Once bacteria reached the correct timepoint (N+ = OD_600nm_, N- = 20 minutes post nitrogen run out as defined previously (9)), 10ml aliquots were removed, bacteria were pelleted and resuspended in 10ml Gutnick minimal media containing 10mM NH_4_Cl supplemented with an antibiotic at a concentration at least 10x MIC (1μg/ml ciprofloxacin or 100μg/ml ampicillin), in 50ml conical tubes. Bacteria were then incubated for 24 hours at 37°C shaking at 180 r.p.m. Samples were taken for serial dilutions and counting of colony forming units (CFU) from the initial cultures (time point 0 hour) and 1, 4 and 24 hours of antibiotic exposure. For CFU counting bacteria were pelleted as 1ml aliquots, the supernatants removed and resuspended in PBS prior to serial dilution and plating and enumeration.

### Determination of minimum inhibitory concentrations (MIC)

Minimum inhibitory concentrations of ciprofloxacin for each strain were determined using an M.I.C. evaluator tests (Oxoid, UK) as described by the manufacturers instructions. Briefly, bacterial cultures were grown overnight in 5 ml cultures either in LB or Gutnick minimal medium containing 10mM NH_4_Cl Bacterial suspensions of 0D600nm = 1 were prepared and 100μl was spread on Iso-sensitest agar plates. An M.I.C. evaluator strip impregnated with an increasing concentration gradient of ciprofloxacin was added to the plate and incubated at 37°C overnight. MIC was determined as the lowest concentration of antibiotic that prevent growth.

### Quantitative Real time PCR (qRT-PCR)

qRT-PCR was completed as described in Brown et al (2014). Briefly, total RNA samples were extracted from cells at specified time-points, RNA was stabilized with Qiagen RNA protect reagent (Qiagen, U.S.A.) and extracted using the PureLink™ RNA Mini kit (Ambion Life Technologies, U.S.A.). Purified RNA was stored at -80°C in nuclease-free water. cDNA was amplified from 100 ng of RNA using the high-capacity cDNA reverse transcription kit (Applied Biosystems, U.S.A.). qRT-PCR reactions were completed in a final volume of 10 μl (1 μl cDNA, 5 μl TaqMan Fast PCR master mix, 0. 5 μl TaqMan probe (both Applied Biosystems, U.S.A.)). Amplification was performed on an Applied Biosystems 7500 Fast Real-Time machine using the following conditions; 50°C 5 min, 95°C 10 min, and 40 cycles of 95°C r sec, 60°C 1 min. Realtime analysis was performed in triplicate on RNA from three independent cultures and quantification of 16S expression served as an internal control. The relative expression ratios were calculated using the delta-delta method (PerkinElmer, U.S.A.). Statistical analysis of data was performed by one-way ANOVA, a *P*-value ≤0.05 was considered to be a significant difference. Primer and probe mixtures were custom designed from Applied Biosystems (Custom TaqMan gene expression Assays (Applied Biosystems, Life Technologies, U.S.A.)) sequences can be viewed in Supplementary Table 1.

## Acknowledgements

This work was supported by an Imperial College Junior Research Fellowship awarded to D.R.B. I would like to thank Gayathri Sritharan for preliminary work to determine the conditions used for the assays and Sivaramesh Wigneshweraraj for discussions and comments on the manuscript.

## References

1. Zimmer DP, Soupene E, Lee HL, Wendisch VF, Khodursky AB, Peter BJ, Bender RA, Kustu S. 2000. Nitrogen regulatory protein C-controlled genes of *Escherichia coli:* scavenging as a defense against nitrogen limitation. Proc Natl Acad Sci USA 97:14674–14679.

2. Reitzer L. 2003. Nitrogen assimilation and global regulation in *Escherichia coli*. Annu Rev Microbiol 57:155–176.

3. Jiang P, Ninfa AJ. 1999. Regulation of Autophosphorylation of *Escherichia coli* Nitrogen Regulator II by the PII Signal Transduction Protein. Journal of Bacteriology 181:1906–1911.

4. Wigneshweraraj S, Bose D, Burrows PC, Joly N, Schumacher J, Rappas M, Pape T, Zhang X, Stockley P, Severinov K, Buck M. 2008. Modus operandi of the bacterial RNA polymerase containing the σ^54^ promoter-specificity factor. Molecular Microbiology 68:538–546.

5. Hartman CE, Samuels DJ, Karls AC. 2016. Modulating *Salmonella* Typhimurium’s Response to a Changing Environment through Bacterial Enhancer-Binding Proteins and the RpoN Regulon. Front Mol Biosci 3:41.

6. Muse WB, Bender RA. 1998. The nac (nitrogen assimilation control) gene from *Escherichia coli*. Journal of Bacteriology 180:1166–1173.

7. Figueira R, Brown DR, Ferreira D, Eldridge MJG, Burchell L, Pan Z, Helaine S, Wigneshweraraj S. 2015. Adaptation to sustained nitrogen starvation by *Escherichia coli* requires the eukaryote-like serine/threonine kinase YeaG. Sci Rep 5:17524.

8. Switzer A, Evangelopoulos D, Figueira R, de Carvalho LPS, Brown DR, Wigneshweraraj S. 2018. A novel regulatory factor affecting the transcription of methionine biosynthesis genes in *Escherichia coli* experiencing sustained nitrogen starvation. Microbiology 65:587.

9. Brown DR, Barton G, Pan Z, Buck M, Wigneshweraraj S. 2014. Nitrogen stress response and stringent response are coupled in *Escherichia coli*. Nat Commun 5:4115.

10. Hauryliuk V, Atkinson GC, Murakami KS, Tenson T, Gerdes K. 2015. Recent functional insights into the role of (p)ppGpp in bacterial physiology. Nat Rev Micro 13:298–309.

11. Nguyen D, Joshi-Datar A, Lepine F, Bauerle E, Olakanmi O, Beer K, McKay G, Siehnel R, Schafhauser J, Wang Y, Britigan BE, Singh PK. 2011. Active starvation responses mediate antibiotic tolerance in biofilms and nutrient-limited bacteria. Science 334:982–986.

12. Helaine S, Cheverton AM, Watson KG, Faure LM, Matthews SA, Holden DW. 2014. Internalization of *Salmonella* by macrophages induces formation of nonreplicating persisters. Science 343:204–208.

13. Verstraeten N, Knapen WJ, Kint CI, Liebens V, Van den Bergh B, Dewachter L, Michiels JE, Fu Q, David CC, Fierro AC, Marchal K, Beirlant J, Versées W, Hofkens J, Jansen M, Fauvart M, Michiels J. 2015. Obg and Membrane Depolarization Are Part of a Microbial Bet-Hedging Strategy that Leads to Antibiotic Tolerance. Molecular Cell 59:9–21.

14. Harms A, Maisonneuve E, Gerdes K. 2016. Mechanisms of bacterial persistence during stress and antibiotic exposure. Science 354:aaf4268.

15. Levin-Reisman I, Ronin I, Gefen O, Braniss I, Shoresh N, Balaban NQ. 2017. Antibiotic tolerance facilitates the evolution of resistance. Science 355:2191–830.

16. Amato SM, Orman MA, Brynildsen MP. 2013. Metabolic Control of Persister Formation in *Escherichia coli*. Molecular Cell 50:475–487.

17. Fung DKC, Chan EWC, Chin ML, Chan RCY. 2010. Delineation of a bacterial starvation stress response network which can mediate antibiotic tolerance development. Antimicrob Agents Chemother 54:1082–1093.

18. Srivatsan A, Wang JD. 2008. Control of bacterial transcription, translation and replication by (p)ppGpp. Current Opinion in Microbiology 11:100–105.

19. Van den Bergh B, Fauvart M, Michiels J. 2017. Formation, physiology, ecology, evolution and clinical importance of bacterial persisters. FEMS Microbiology Reviews 41:219–251.

20. Schumacher J, Behrends V, Pan Z, Brown DR, Heydenreich F, Lewis MR, Bennett MH, Razzaghi B, Komorowski M, Barahona M, Stumpf MPH, Wigneshweraraj S, Bundy JG, Buck M. 2013. Nitrogen and carbon status are integrated at the transcriptional level by the nitrogen regulator NtrC in vivo. MBio 4:e00881-13.

21. Girard ME, Gopalkrishnan S, Grace ED, Halliday JA, Gourse RL, Herman C. 2017. DksA and ppGpp Regulate the σ^S^ Stress Response by Activating Promoters for the Small RNA DsrA and the Anti-Adapter Protein IraP. Journal of Bacteriology 200:e00463-17.

22. Hansen S, Lewis K, Vulić M. 2008. Role of global regulators and nucleotide metabolism in antibiotic tolerance in *Escherichia coli*. Antimicrob Agents Chemother 52:2718–2726.

23. Stewart PS, Franklin MJ, Williamson KS, Folsom JP, Boegli L, James GA. 2015. Contribution of stress responses to antibiotic tolerance in Pseudomonas aeruginosa biofilms. Antimicrob Agents Chemother 59:3838–3847.

24. Hong SH, Wang X, O’Connor HF, Benedik MJ, Wood TK. 2012. Bacterial persistence increases as environmental fitness decreases. Microb Biotechnol 5:509–522.

25. Wu N, He L, Cui P, Wang W, Yuan Y, Liu S, Xu T, Zhang S, Wu J, Zhang W, Zhang Y. 2015. Ranking of persister genes in the same Escherichia coli genetic background demonstrates varying importance of individual persister genes in tolerance to different antibiotics. Front Microbiol 6:1003.

26. Radzikowski JL, Vedelaar S, Siegel D, Ortega ÁD, Schmidt A, Heinemann M. 2016. Bacterial persistence is an active σ^S^ stress response to metabolic flux limitation. Mol Syst Biol 12:882.

27. Baba T, Ara T, Hasegawa M, Takai Y, Okumura Y, Baba M, Datsenko KA, Tomita M, Wanner BL, Mori H. 2006. Construction of *Escherichia coli* K-12 in-frame, single-gene knockout mutants: the Keio collection. Mol Syst Biol 2:2006-0008.

28. Wang B, Kitney RI, Joly N, Buck M. 2011. Engineering modular and orthogonal genetic logic gates for robust digital-like synthetic biology. Nat Commun 2:508.

29. Atlas RM. 2010. Handbook of microbiological media, vol 1. Fourth edition. CRC Press.

30. Schneider DA, Murray HD, Gourse RL. 2003. Measuring control of transcription initiation by changing concentrations of nucleotides and their derivatives. Meth Enzymol 370:606–617.

